# Probing Local Force Propagation in Tensed Fibrous Gels

**DOI:** 10.1101/2022.09.07.506942

**Authors:** Shahar Goren, Maayan Levin, Guy Brand, Ayelet Lesman, Raya Sorkin

## Abstract

Fibrous hydrogels are a key component of soft animal tissues. They support cellular functions and facilitate efficient mechanical communication between cells. Due to their nonlinear mechanical properties, fibrous materials display non-trivial force propagation at the microscale, that is enhanced compared to that of linear-elastic materials. In the body, tissues are constantly subjected to external loads that tense or compress them, modifying their micro-mechanical properties into an anisotropic state. However, it is unknown how force propagation is modified by this isotropic-to-anisotropic transition. Here, we directly measure force propagation in tensed fibrin hydrogels. Local perturbations are induced by oscillating microspheres using optical tweezers. We use both 1-point and 2-point microrheology to simultaneously measure both the shear modulus and force propagation. We suggest a mathematical framework to quantify anisotropic force propagation trends. We show that force propagation becomes anisotropic in tensed gels, with, surprisingly, stronger response to perturbations perpendicular to the axis of tension. Our results suggest that under external loads, there are favoured directions of mechanical communication between cells in a tissue. Importantly, we also find that external tension increases the range of force transmission by altering the power-law exponent governing the decay of oscillations with distance from the perturbation. We end with a discussion of possible implications and future directions for research.

## 1 Introduction

Soft animal tissues are structured by the extracellular matrix (ECM) [1], a branched network of fibers, to which cells adhere. The fibrous materials that compose the ECM have remarkable nonlinear properties [2, 3] – for example, unlike most synthetic materials, the ECM can undergo very large deformations and recover without damage due to strain stiffening [4]. Cells can modify the ECM by applying local forces [5,6], and also respond to the ECM mechanical properties [7–9]. Due to its role in many biological processes [8,10–12], the mechanical properties of the ECM has been extensively studied.

Over the last decades, it was found that during processes such as blood vessel growth [13], wound healing [14] and cancer metastasis [15, 16], cells communicate their position and orientation, and sense other nearby cells or rigid boundaries. They do so by applying contractile forces and sensing the mechanical environment. Hence, mechanical signals emerged as a well-recognized type of cellular communication. The nonlinear mechanics of the ECM was shown to facilitate and enhance mechanical communication between cells [17–19]. Nonlinear materials can exhibit non-trivial, longer-ranged force propagation properties, that facilitate efficient cell-cell interactions [20–24]. In an organism, fibrous materials are constantly subject to external and internal stresses due to body posture and movement, blood flow and muscle contraction [25]. External tension affects various cellular processes. For example, it affects cell migration in wound closure [26,27], and can facilitate muscle [28] and blood vessel [29] development. These effects are often attributed to the nonlinear mechanical response of the underlying scaffold to external tension. Tension can cause fibrous materials to align and stiffen, and thus direct cell behavior. However, it is unknown how these tension-induced alignment and stiffening effects are manifested in the microscale mechanics that cells feel, and in particular, their effect on the propagation of mechanical signals is unknown. We have recently shown that the anisotropic state of the material under tension is a critical parameter that determines the capacity of fibrous networks to transmit displacements away from their source [23], motivating a closer look at the anisotropy induced by tension and its relationship to force propagation, and ultimately to cell-cell communication.

Bulk shear rheology using a rheometer is the most prominent method to characterize the nonlinear mechanical properties of fibrous materials. [2,30–34]. Bulk rheology studies have found that all biological fibrous materials stiffen under shear and tension [2], and that under compression, there are both softening and stiffening regimes [30]. The leading suggested mechanisms that may account for this rich behavior are the alignment of fibers under tension [18, 35–37], buckling of fibers under compression [38–40], and bending-to-stretching rigidity transition arising from the low-connectivity topology of these networks [41,42]. However, despite the ability of rheometers to probe the nonlinearity of fibrous materails, they can only do so at the macroscopic scale of the gel as a whole, and do not provide direct information on the mechanics at the microscale. Thus, despite accumulated knowledge on the macro-mechanical and biological consequences of external tension, there is not yet a good understanding of how tension affects the mechanical micro-environment in which cells function. In addition, bulk rheology does not provide any direct information on the propagation of mechanical signals.

A more recent approach to the study of mechanical properties of fibrous materials is microrheology. Microrheology allows inferring the viscoelastic properties of a material from the movement of micro-particles embedded in it [43]. Microrheological techniques are categorized as either passive or active, depending on whether thermal fluctuations or active external forces drive the particles [44]. Yet another distinction is between 1-particle methods, which track only single-particle displacements, and 2-particle methods, in which correlations between pairs of particles are used to determine the medium viscoelastic properties. Microrheology readily provides microscopic mechanical viscoelastic information. Also, microrheology is a natural tool to study the propagation of mechanical signals due to its access to local forces and displacements. However, it is usually used to study the linear response regime, and does not provide access to the nonlinear response, which is only triggered at large deformations.

Previous microrheology studies have focused on characterizing the mechanics of biological gels such as collagen, fibrin and actin under different preparation conditions [45–47] or during different stages of their formation [48–50]. Others have also used 1-particle microrheology to show how local stiffness of fibrous gels is modified by cell contractility [51, 52] or internal shear [53, 54]. Few works have also fabricated and characterized anisotropic fibrous gels [55, 56]. However, none of these studies quantitatively characterized force transmission in the deformed environment.

Here we develop a novel experimental technique to probe the microscale nonlinear mechanical properties of fibrin gels. Fibrin, being the main structural component of blood clots [33, 57, 58], is a relevant model for wound healing environments, in which cells need to communicate in order to orchestrate the healing reaction. We combine a method to apply controlled stretch to soft fibrous gels with optical tweezers active microrheology. Our approach is to use external tension to trigger a nonlinear mechanical response in the material, leading to a mechanically anisotropic microenvironment. We then use microrheology together with fluorescent confocal microscopy to characterize the modified microenvironment and force transmission. We quantitatively analyze force transmission in fibrin gels under tension, providing a framework for characterizing anisotropic materials’ response to local perturbations. Our goal is to explore the relationship between tension, fiber alignment and force transmission, improve the understanding of the role of external tension in biological processes, and promote the development of better models of fibrous materials. Overall, we show that tension leads to anisotropy in the local stiffness and in force transmission. We observe enhanced material response in tensed materials, and discuss possible causes and implications.

## 2 Methods

### 2.1 Fibrin gel preparation

Fibrin gel is prepared by mixing Fibrinogen (Evicel Biopharmaceuticals) at a final concentration of 0.25 mg/ml with Thrombin (Evicel Biopharmaceuticals) at a final concentration of 1 U/ml. The fibrinogen is fluorescently labeled with Alexa Fluor 546, succinimidyl ester (Invitrogen), as previously described [59]. For isotropic gels, the solution is mixed on top of a glass slide, and closed with another slide with 1mm spacing. Gels that will be stretched are prepared on top of a parafilm strip, inside a circular mold cut into a silicone rubber strip of 1mm thickness, as we described previously [59, 60]. The gels polymerize over-night at 37°c.

### 2.2 Bulk Rheology

We use a Discovery HR-3 rheometer with a plate-plate geometry (20 mm diameter, 1mm gap) to characterize the macroscopic shear modulus of the gel at increasing degrees of strain. The gels are polymerized inside the rheometer for 3 hours before measurements. We choose a strain frequency of 0.5 Hz, matching the frequency in our microrheology experiments.

### 2.3 Passive Microrheology

Imaging the motion of the tracer particles within the gels was done using an Olympus IX71 epifluorescence microscope, at λ = 480 nm with a 60× oil objective. We recorded the motion of approximately 250 particles in the field of view using a CMOS video camera (Grasshoper 3, Point Gray) at a frame rate of 30 Hz with an exposure time of 10 ms. We used data from at least 105 frames. Particle tracking was done by conventional video microscopy, using the protocol of Crocker and Grier (59). The mean square displacement and *G’, G”* were computed using previously described methods (47, 60).

### 2.4 Loading of gels into tensile device

The tensile device used in this work was a simple metal frame with clamps on both sides and a holder for a glass slide in the middle (see supplementray material, Figure S1). The silicon strip including the gel is picked up and the parafilm is removed. The silicone strip is then placed along the frame so that the gel is laying on the glass slide and the ends of the strip are under the clamps. The silicone strip is then clamped on one side, pulled manually from the other side so that the length of the hole enclosing the gel in increased by 50%. The other clamp is then closed, preventing the silicone from contracting back. The frame is then placed on the stage of the optical tweezers - confocal fluorescence microscope (C-trap; Lumicks). Within this setup, the axis of tension is always chosen as the x-axis of the microscope imaging. For isotropic gels, samples were prepared on a glass cover-slip, placed on the microscope stage and held with clamps.

### 2.5 Force calibration

The trap stiffness *κ* was calibrated by fitting the power spectrum of beads trapped in water to a lorenzian [61]. Calibration was done separately for each laser power used in the experiment.

### 2.6 Active Microrheology

We use a C-Trap ® confocal fluorescence optical tweezers setup (LUMICKS) made of an inverted microscope based on a water-immersion objective (NA 1.2) together with a condenser top lens. The optical trap is generated by 10W 1064-nm laser. The trap can be steered using a piezo mirror with a feedback mechanism to determine its accurate position. The samples were illuminated by a bright field 850-nm LED and imaged in transmission onto a metal-oxide semiconductor (CMOS) camera at a frame rate of 15 Hz. Each microrheology measurement consists of focusing the trapping laser on a 3.5 micron bead, and driving it sinusoidally by oscillating the trapping laser once in the x-axis and once in the y-axis. The drive frequency was chosen as 0.5 Hz. The drive amplitude was 1-2 microns (but was always equal in x and y), and the duration of each oscillation was 10-20 seconds. The driven bead was tracked using phase-modulated Hough transform with Matlab (The MathWorks, Inc.). The same method also yields the bead radius *a* which was used in computations to prevent errors from bead size dispersity. The material shear modulus *G* was estimated using Stokes’ law, linking the displacement 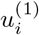 of a bead of radius *a* subject to a force *F_i_*:

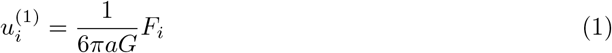

with the force given by 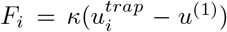. Simultaneously, tracer beads were tracked using the trackpy python package [62]. The oscillation amplitudes of driven beads and tracer beads on both axes were found by taking the Fourier transform of their trajectory along the desired coordinates, and extracting its magnitude at the applied frequency. This effectively filters out low-frequency drifts and high-frequency noise, leaving only the response to the applied perturbation. The accuracy of this method was tested by oscillating a 1.5 micron bead in water, as described in the supplementary information, Figure S2.

### 2.7 2-point active microrheology analysis

In order to suggest a framework for analyzing 2-particle microrheology in tensed gels, we first discuss the suitable theory for isotropic fibrin gels. [63, 64] When a bead is driven within a linear, isotropic medium by optical tweezers, the displacement of a second bead at position 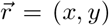 with respect to the driven bead is given by the generalized Oseen’s tensor:

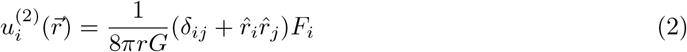

With 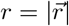 and *i,j* ∈ (*x,y*). Instead of this expression, we use an angle-integrated expression for 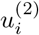 in terms of 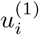 (derived in the supplementary information, section 1):

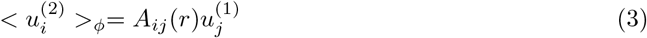

Where *ϕ* = *tan*^-1^ (*y*/*x*). For an isotropic medium, equations 1 and 2 yield:

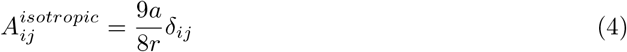

While it is customary to choose a coordinate system in which the 2 particles are aligned along the x axis, here we deal with anisotropic materials which have fixed principal directions. Throughout this paper, *x* is chosen as the axis along which the gel is tensed. For an isotropic material, the off-diagonal elements of *A_ij_* vanish due to inversion symmetries *x* ⇔ −*x* and *y* ⇔ −*y*. These symmetries are expected to hold also for tensed gels, because the chosen coordinate system is aligned with the principal axes of the material. However, the 2 diagonal elements will equal only in isotropic gels, due to the rotational symmetry. This symmetry breaks when the medium is subject to tension along the *x* or *y* directions. We can quantify the anisotropy by measuring the diagonal elements of *A_ij_*(*r*). The response tensor *A_ij_*(*r*) was computed by collecting the positions and amplitudes of tracer beads from multiple positions and gels, normalizing them by the driven bead amplitude, and averaging over their values in concentric rings. This method was necessary in order to overcome the material heterogeneity, thus requiring us to average over large sets of measurements to characterize the material response. For quantitative analysis, data was collected with 10-30 driven beads in each gel, and for 16 tensed and 16 untensed gels.

### 2.8 Fluorescent microscopy

For each microrheological video, we also acquire a fluorescent scan of the structure of the fiber network in that region. 2 excitation lasers, of 488nm and 561nm, to excite the tracer beads and labeled fibers respectively. Scanning was done using a fast tip/tilt piezo mirror. Photons were counted using fiber-coupled single-photon counting modules. The multimode fibers serve as pinholes providing background rejection. This enables us to estimate the geometrical anisotropy in the surrounding material, manifested by fiber alignment in the direction of the stretched axis.

### 2.9 Quantification of alignment

We use Matlab to estimate the alignment of fibers from confocal images. The green fluorescent label of the fibers are used for this analysis. Local intensity gradients *g_x_, g_y_* were calculated for each pixel by convolution of the image with 21X21 Sobel operator, and the fiber orientation was calculated as *θ* = *tan*^-1^ (*gx/gy*). The nematic order paremener (NOP) was calculated as:

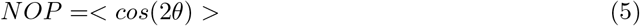

The nematic order parameter varies between −1 and 1, with 1 corresponding to full alignment in the x direction and 0 corresponding to random orientation.

### 2.10 Statistics

To determine whether 2 measured distributions (for example, shear modulus distributions) are distinguishable, we used Kolmogorov-Smirnov test. This test is suitable because it makes no previous assumptions about the underlying distributions. In all cases, we denote a P-value (*P_KS_*) less than 10^-4^ with ****.

## 3 Results

### 3.1 Fibrin gels are predominantly elastic

We first perform bulk rheology measurments to assert that fibrin gels are solid under the preparation conditions chosen, and characterize their macroscopic linear and nonlinear shear modulus. As can be seen in Figure 1A, we find characteristic stiffness of approximately 0.7 Pa, with solidlike viscoelastic behavior (*G*′ > *G*″). At strains of approximately 20%, nonlinear strain-stiffening begins, in line with previous findings [2]. Here, stiffness increases by about a factor of 4 at shear strains of around 100%. We hypothesize that this strain-stiffening behavior is related to fiber alignment and manifested at the microscale by anisotropic mechanics, which can be determined from microrheology. Figure 1B plots the frequency dependence of the shear modulus, for bulk rheology (for 5% strain) and and 1- and 2-particle microrheology of isotropic gels. For all 3 methods, we observe a crossover between *G*′ and *G*″ as was found in other studies [65]. Below the crossover, we observe a plateau in *G*′, corresponding to a stable gel regime. All 3 methods give similar values of *G*′ for low frequencies. The crossover between *G*′ and *G*″ curves also occurs at a slightly different frequency for each method, meaning that their relative sensitivity to the elastic and viscous components is somewhat different. The fact that 1-particle and 2-particle methods yield similar values of *G*′ at low frequencies is not trivial [64], and implies that the beads are strongly coupled to the fibers, so single bead thermal fluctuations are sensitive to the gel mechanics rather than to local interactions between the fibers and solvent. Another important observation is that compared to bulk rheology, G” is negligible compared to G’ in passive microrheology for low frequencies. This once again implies that the solvent contribution to the bead dynamics at frequencies below 10Hz is negligible. For low frequencies (below 10Hz), gels can be considered as an elastic solid that is merely stabilized by the solvent, but whose mechanics are dominated by the elastic network, as far as bead dynamics are concerned. We will thus focus on low frequencies (frequency *f* = 0.5*Hz*, angular frequency *ω* ≈ 3Hz) and assume *G* = *G*′ with negligible frequency dependence.

**Figure 1:**
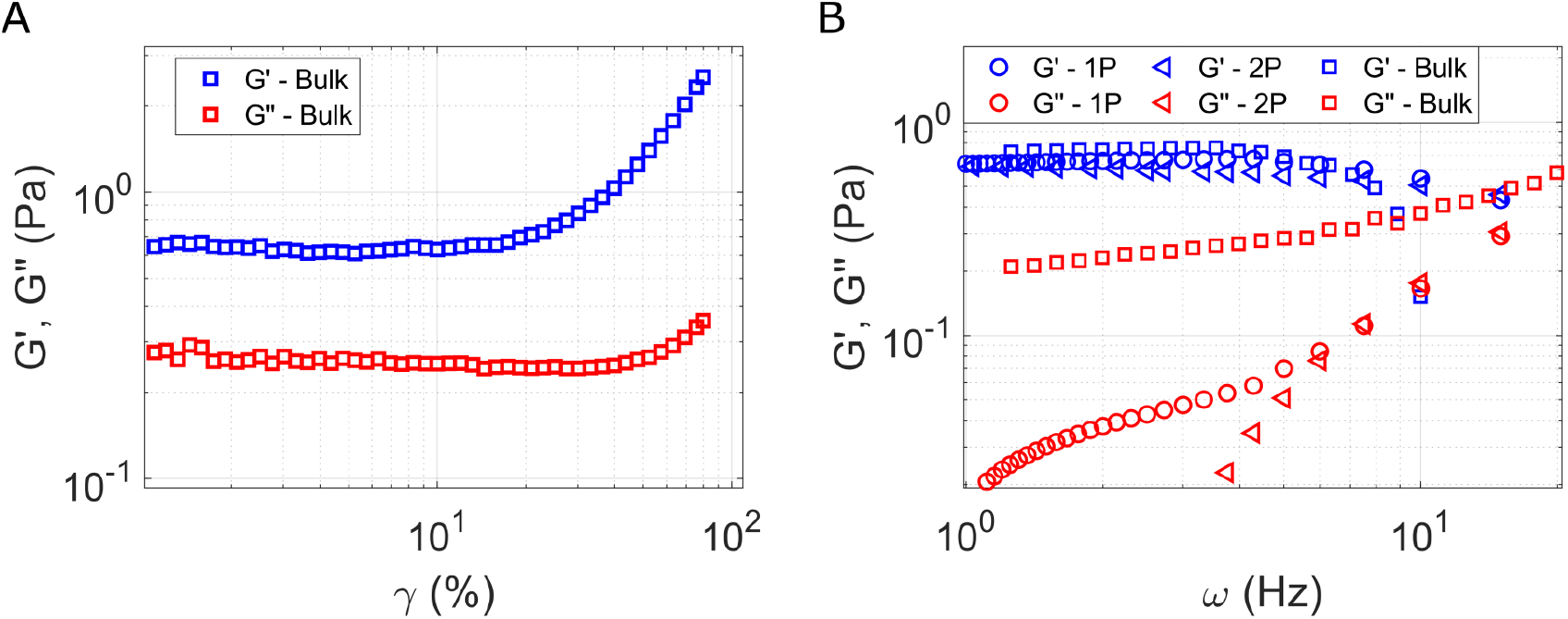
Bulk rheology and passive microrheology. (A) The viscoelastic modulus of fibrin under increasing shear strain, showing characteristic stiffening. Frequency was 0.5 Hz. (B) Frequency dependence of the viscoelastic modulus from bulk rheology, 1-particle and 2-particle passive microrheology.

### 3.2 Tension-induced transition from isotropic to anisotropic state

Fluorescent microscopy shows substantial fiber alignment in the *x* direction in tensed gels, and isotropic distribution of fiber orientations in untensed gels, as seen in Figure 2A,B. Stretching the fibrous network also leads to contraction in the *y* and *z* directions, and therefore the overall density of fibers in the field of view is higher for tensed gels. Figures 2C,D show the distribution of orientations from which the nematic order parameter is computed. Tension leads to a large fraction of fibers aligned in the x direction, and a peak is observed in *θ* = 0 for tensed gels. Confocal microscopy has a directional axis of scanning, which introduces spurious directionality in confocal scans. Most images in this study where recorded with *x* as the fast axis, and as can be seen in the figure 2C, a small peak appears for untensed gels as well. We correct for this effect by quantifying and subtracting it as described in the Supplementary Information Figure S3. Figure 2E compares the distribution of the NOP for untensed and tensed gels, demonstrating the transition from isotropic to anisotropic state due to tension.

**Figure 2:**
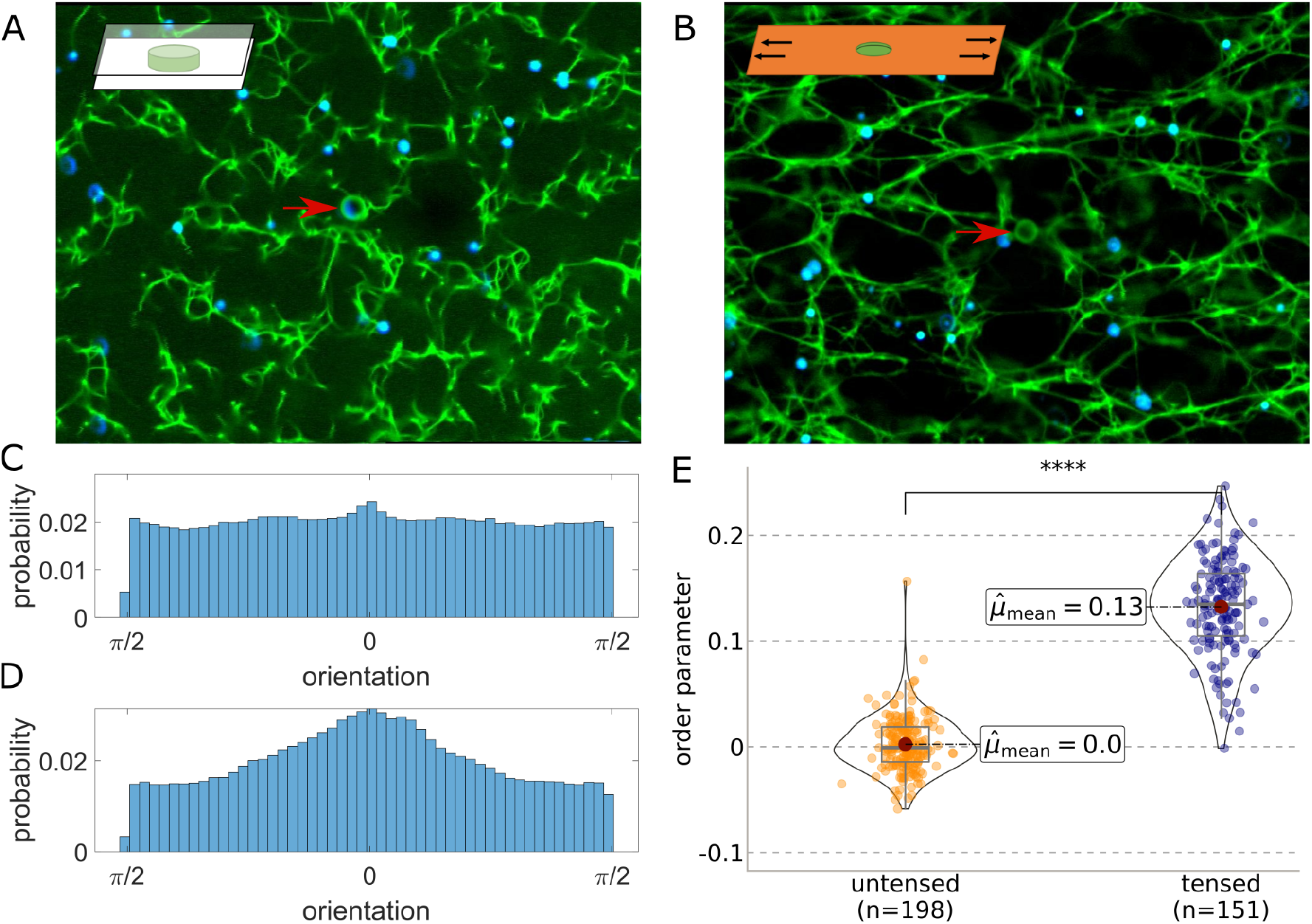
Confocal fluorescent microscopy shows fiber alignment in tensed gels. (A) Untensed and (B) Tensed gels confocal scans. Driven bead is pointed with an arrow in the center, and tracer beads are shown in blue. Insets describe the sample loading conditions for each type. (C) and (D) display the orientation distribution for untensed and tensed gels, respectively. (E) Comparison of the distributions of order parameter *NOP* =< *cos*(2*θ*) > computed over the directions of intensity gradients in image scans in untensed and tensed gels.

### 3.3 Tension-induced stiffness anisotropy measured by 1-point microrheology

The embedded microsphere dynamics are coupled to those of the fibrin network, which is solid under the experimental conditions. This is due to the following reasons: first, despite its low concentration, fibrin still forms a branched network that is stable under the forces applied in the experiment. Second, the beads in the experiment adhere strongly to the fibers (as can be seen in the confocal images in Figure 2), and are therefore coupled to the network at all relevant timescales, and are less sensitive to the solute viscosity at low frequencies [66]. Lastly, our experiments focus on the low frequency regime (~ 1*Hz*), in which the medium is predominantly elastic, as seen by bulk rheology (Figure 1). Thus, the measured bead dynamics reflect the elastic shear modulus of the gel. To simplify our analysis, we also assume that the network is incompressible (with Poisson ratio *ν* = 1/2). This assumption may be inaccurate, but the errors it introduces are in the order of several percent at most [67]. We compare compressible and incompressible expressions in the Supplementary Information, section 1.

We quantify the modification in stiffness and mechanical anisotropy as a result of external tension. First, to observe how the fibrous network deforms in response to the oscillation of the driven bead, we performed a step-by-step oscillation experiment, in which we translated the driven bead in small consecutive steps composing an oscillation, while taking a confocal scan beween each 2 consecutive steps. Video 1 displays this process, showing the driven bead as a dark spot in the middle, the fibrin fibers in green and the tracer beads in orange. In practice, however, we use brightfield videos, such as Videos 2 and 3 in order to track the driven bead, compute its amplitude and obtain the local shear modulus from equation 1. Figures 3 A and B compare the distributions of shear moduli in *x* and *y* directions for untensed gels and tensed gels, respectively. It can be seen that for untensed gels, stiffness values of ~ 0.44 ± 0.36*Pa* were found. It can be seen that untensed gels are isotropic, with their distributions for *x* and *y* shear moduli are indistinguishable (Kolmogorov-Smirnov, *P_KS_* = 0.44). For tensed gels we observe distinctly different distributions for *x* and *y* (*P_KS_* < 10^-4^) directions with moduli of ~ 16 ± 14*Pa* in the direction of tension (*x*) and ~ 9 ± 8*Pa* in the perpendicular direction (*y*). This corresponds to a remarkable stiffening of 20-40 fold compared to untensed gels. The mechanical anisotropy, i.e. the ratio *G_y_/G_x_* decreases from 1 in untensed gels to ~ 0.62 ± 0.27 on average in tensed gels (Figure 3C). Our results demonstrate that tension leads to a dramatic increase in stiffness in all directions, and a moderate anisotropic effect. Figure 3D plots the mechanical anisotropy ratio *G_y_/G_x_* and its dependency on the local fiber alignment in the vicinity of the driven bead. It can be seen that fiber alignment is a predictor of mechanical anisotropy, for both untensed and tensed gels.

**Figure 3:**
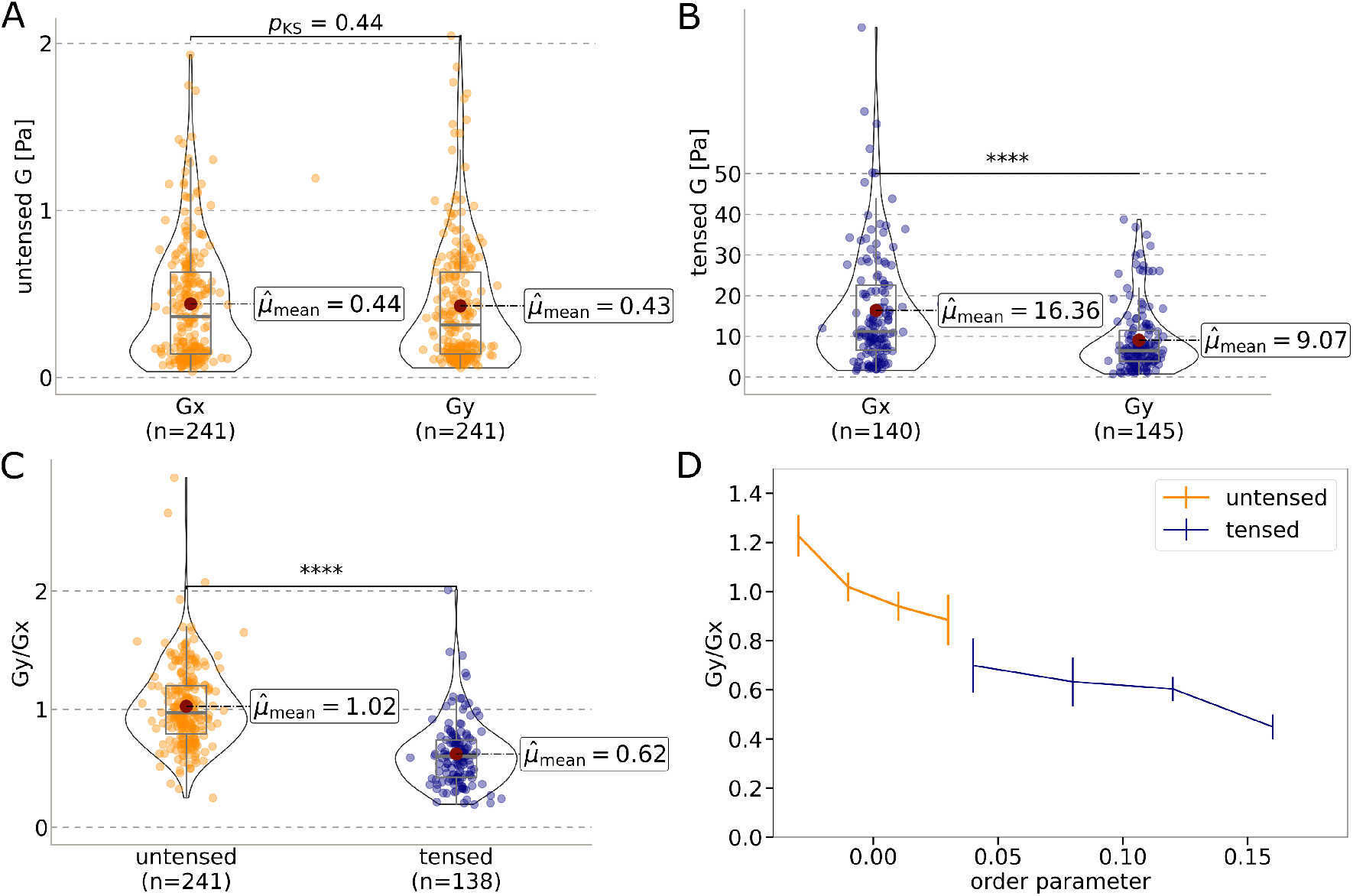
Local stiffness measurement of fibrin gels with 1-point microrheology. (A) and (B) display comparison of the shear modulus distributions in x and y directions, for untensed and tensed gels respectively. (C) Distributions of stiffness ratios *G_y_/G_x_*, in untensed and tensed gels. Horizontal black lines represent Kolmogorov-Smirnov statistical tests, where **** corresponds to a p-value lower than 0.0001. (D) Stiffness ratio as a function of the order parameter measured from a confocal scans in the driven bead area. Trends display average ± SEM of binned distributions.

### 3.4 Force transmission of untensed and tensed gels measured with 2-point microrheology

2-point microrheology experiments are performed by oscillating one bead and measuring the response amplitude of other beads in its vicinity. A crucial assumption in our analysis is that the response to the perturbation is linear, i.e. the oscillation amplitudes of each of the beads in the field of view is linearly proportional to the amplitude of the driven bead. We assume this is the case for both untensed and tensed gels. That is, the external tension triggers a nonlinear material response, and the small bead oscillations probe the modified environment after the response to tension, without an additional nonlinear effect. We check this assumption in Figure S4 in the supplementary material. We also assume that for low frequencies the response is independent of the frequency, as described above. Both these assumptions are justified as discussed and verified in the supplementary material, Figure S5.

Videos 2 and 3 display driven bead oscillations in the *x* and *y* directions, showing also the tracer beads’ response. Figure 4 displays a typical example of quantification of tracer bead amplitudes, directions and phases of oscillations in *x* and *y* for an isotropic, untensed gel. The measured oscillations correspond roughly to Oseen’s tensor structure, with larger amplitudes along the axis of perturbation. The phase differences obtained from Fourier analysis (Figure 4 C and D) are distributed narrowly around 0, and are probably due to measurement errors rather than actual phase difference. The ratio of drive frequency to frame rate gives 12° phase difference per frame, so that errors around 12° are expected. These results also demonstrate that in the absence of external tension, fibrin gels behave as predicted by Oseen’s tensor, without any time-lag between the oscillations in different regions.

**Figure 4:**
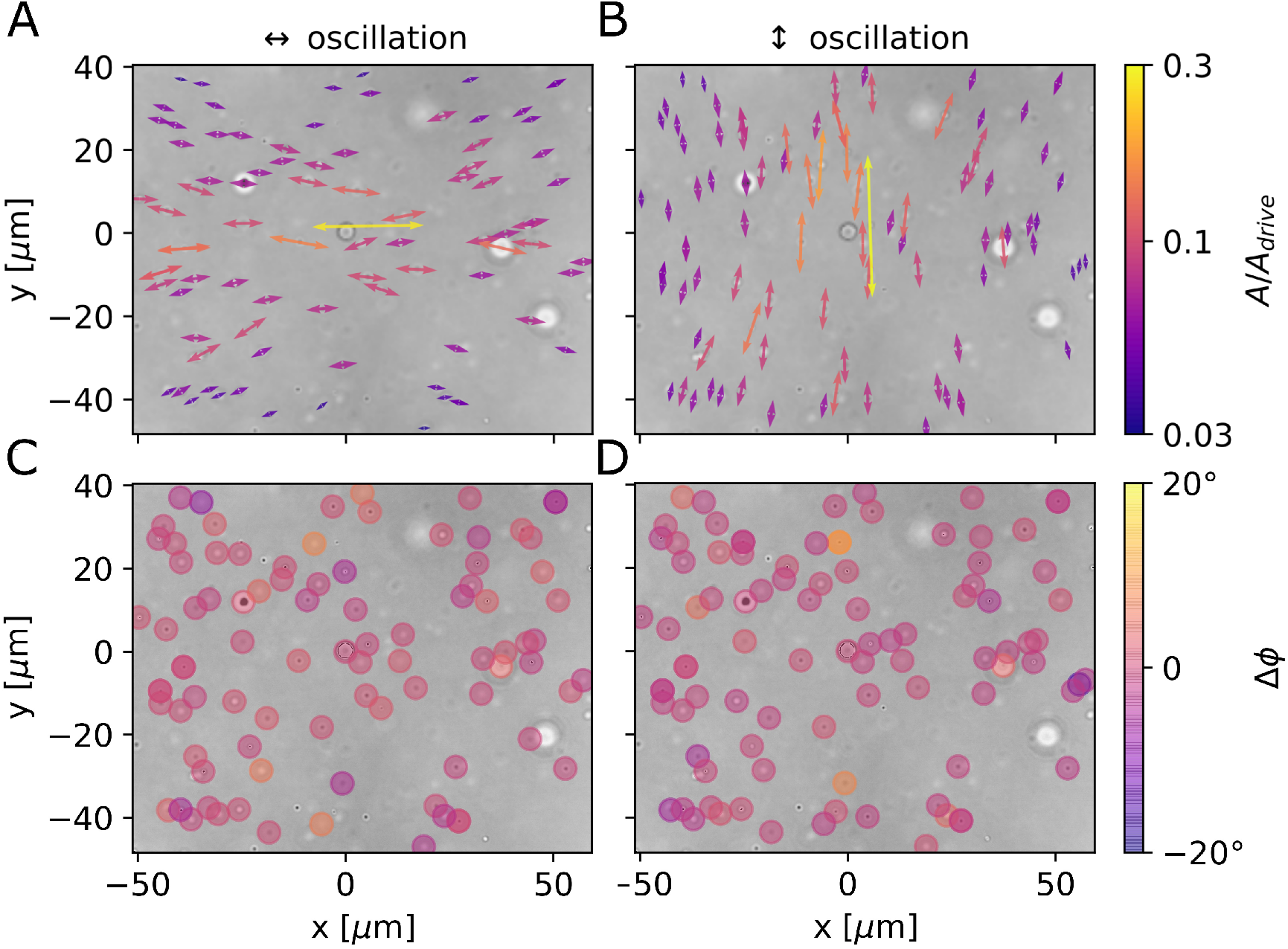
Example of an oscillation field obtained from analysis of a brightfield video in an untensed gel, (A) displaying the response to an oscillation in the x direction, and (B) to oscillation in the y direction. Arrow amplitudes are scaled up by a factor of 10, their direction represents the true oscillation direction, and their color represents their amplitude normalized by the driven bead amplitude. (C) and (D) plot the phase differences Δ*ϕ* = *ϕ_tracer_* – *ϕ_driven_* corresponding to the oscillations in (A) and (B).

To get a reliable quantification of the response tensor, normalized amplitudes of tracer beads from multiple videos are collected. Assuming a linear response of the medium, normalized amplitudes from different videos can be treated as independent measurements of the medium response, which can thus be averaged.

Figure 5 shows collected amplitudes from 10-30 regions in each gel, from 16 untensed (Figure 5 A and B) and 16 tensed (Figure 5 C and D) gels in total. We separate the response for x and y oscillations. We first make sure isotropic gels have identical response in x and y, and then check how this changes for tensed gels.

**Figure 5:**
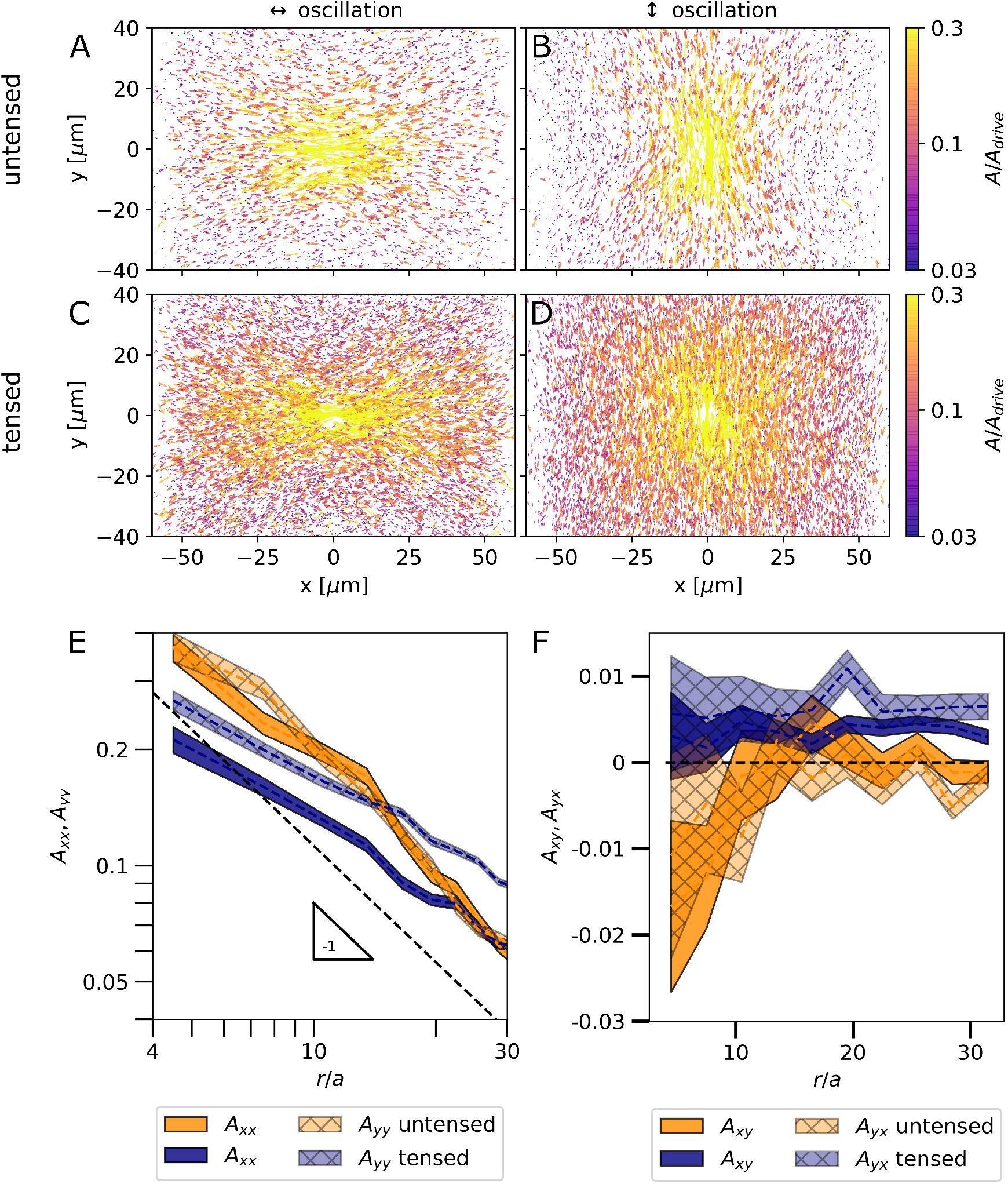
Oscillation fields collected from multiple gels and positions. (A)-(D) display response to oscillations in x and y directions, for untensed and tensed gels. (E) plots the diagonal elements of the response tensor. (F) plots the off-diagonal elements of the response tensor. Trends display average ± SEM

Figure 5 E,F shows the components of the response tensor *A_ij_* as a function of normalized distance from the driven bead (*r/a*, where *a* is the driven bead radius), for tensed and untensed gels. In Figure 5E the diagonal elements *A_xx_* and *A_yy_* are displayed. Several important observations can be made: for untensed gels, the diagonal components of the response tensor are equal and decrease as 1/r, as expected. However, the prefactor deviates from the value of 9/8 predicted by Oseen’s tensor. On average, we measure a prefactor of 1.85, that is about 65% higher than the Oseen’s prefactor, with minor differences between individual gels (see supplementary information, Figure S6). This discrepancy is most likely a result of a small difference between 1-particle and 2-particle measures of the elastic modulus. Microrheological measurements of the shear modulus often differ between 1-point quantifications (from equation 1) and 2-point quantifications (from equation 2). Such discrepency can introduce a correction factor to *A_ij_*.

For tensed gels, *A_yy_* is larger than *A_xx_* by roughly 50%, showing a substantial anisotropic response. In essence, this result means that when the gel is stretched along the x-axis, the transmission of transverse perturbations (in the y-axis) is enhanced with respect to transmission of parallel perturbations. Also, it can be seen in Figure 5E that for tensed gels, the perturbation propagates with power dependence that is smaller than the expected −1. This irregular behavior is consistent in all tensed gels, as shown in the supplementary information, Figure S7. It is remarkable because it cannot be simply explained by anisotropy, and we discuss possible reasons for this behaviour in the discussion section. Looking at the off-diagonal elements (Figure 5F) we can see that for both tensed and untensed gels they are negligibly small, as predicted from symmetry (see section 2.7), although for tensed gels there is a more substantial non-zero off diagonal response. This can be due to a minor misalignment between the tension axis and the optical *x* axis. To demonstrate that our results are robust, we compare the response tensor for 3tensed and 3 untensed gels (supplementary information, figures S6 and S7). We chose gels for which enough data was collected in order to quantify the response tensor without further averaging.

### 3.5 Discussion

Here, we have developed a framework for quantitative analysis of 1-point and 2-point microrheology of fibrous materials under tension. This is the first study, to our knowledge, that demonstrates the use of quantitative microrheology methods, and in particular 2-particle microrheology, for the study of fibrous gels under external tension. Therefore, it has the potential to push forward the understanding of mechanical communication between cells, which plays a pivotal role in many fundamental biological processes including cell migration and proliferation [13, 14, 16, 68, 69]. As most of our body tissues are constantly subject to external loads, it is of great importance to study the propagation of mechanical signals in tissues and tissue-mimicking gels under tension, as we do here.

We combine 3 main approaches to quantify the changes induced by tension – confocal microscopy, 1-point microrheology and 2 point microrheology. Confocal microscopy visualizes the structure of the gel in thin slices. It can be seen in the confocal images that the gels are very sparse, with large pores and fiber lengths with sizes comparable to the diameter of driven beads (~ 3.4*μm*). When tensed, however, a larger fraction of the fibers aligns with the focal plane and the networks appear somewhat denser, with some fibers spanning more than 10 microns along the axis of tension. We provide a simple, robust method to estimate this fiber alignment quantitatively and remove experimental artifacts. We show that our method indeed distinguishes between populations of tensed and untensed gels by their fiber alignment extent.

1-point microrheology has the advantage that it can quantify the nonlinear stiffening induced by external tension. We first compared 1-point microrheology of untensed gels to bulk rheology and passive microrheology and found an acceptable agreement, although active microrheology seems to be somewhat biased towards lower values of *G*. Remarkably, we observe very dramatic stiffening of up to 40-fold increase in stiffness for tensed gels, combined with a x-y stiffness anisotropy. The dramatic stiffening probably arises from multiple mechanisms: it likely involves densification of the gel in the y and z directions, known to give rise to strong stiffening at large deformations [57]. Individual fiber nonlinearity likely contributes to stiffening as well. Fiber nonlinearity and fiber alignment can both contribute to the observed stiffness anisotropy [23].

We used untensed fibrin gels as a control to test our setup and analysis. As expected, we found an isotropic response, with equal diagonal entries of the response tensor, and a characteristic 1/r decay trend. However, there was a small discrepancy between the measured prefactor and the prefactor predicted by Oseen’s tensor. This discrepancy cannot be explained by Poisson’s effect, because compressible corrections are always negative [67], unlike the positive correction we observe. Also, in equation 4, we assumed that the value of the shear modulus obtained from 2-particle microrheology agrees with that of 1-particle microrheology. If they do not completely agree, it can explain the observed discrepancy.

The study of anisotropic materials, and especially their micromechanical properties, is very limited compared to the understanding of isotropic media. The simplest anisotropic material, known as transverse-isotropic material, has 5 elastic moduli [70]. A few studies addressed 1-point microrheology [71] and 2-point microrheology [72] in transverse-isotropic materials, and given the material macroscopic moduli, its response can be predicted. However, since there is no established experimental scheme to measure the moduli of tensed gels, direct comparison of our results with these models is prohibited. Computational and theoretical studies may fill this gap in the future.

Our most important findings result from the quantification of mechanical force propagation in fibrous gels under tension. We hypothesized that the renowned stiffening of fibrin gels should result in anisotropic transmission of mechanical signals. We chose to focus on the angle-averaged response to perturbations in parallel and perpendicular to the axis of tension, and found that the response of the material to oscillations perpendicular to the axis of tension (*y*) is stronger than the response to parallel oscillations. That is, not only is it easier to pull or push along the perpendicular direction to tension axis (*G_y_* < *G_x_*), the material also responds more sensitively to perturbations along this axis. This indicates that tension can give rise to preferred directions of communication between cells, such as during blood vessel growth [13]. Interestingly, we observe an irregular response of tensed gels, in which oscillations decay with an apparent power-law *r^n^* with a decreased exponent *n* > −1, resulting in a longer range. This result motivates future investigations, because the expected power-law *r*^-1^ in the far region arises from material homogeneity (uniform *G*) assumption and the equations of elasticity. Since our results are averaged over many regions, inhomogeneity is not expected to lead to robust power-law other than *r*^-1^ while we show that it is indeed robust, see supplementary information section 7. The usual cause for irregular power-law behavior is a non-linear response, but we also ruled this out by measuring a linear response to different amplitudes (see supplementary information section 5 and Figure S4). We suggest that tension deforms the gel structure, changing its typical length scales. If a material length scale (for example, the pore length along the *x* direction) increases and becomes large compared to the field of view, it is possible that the far field *r*^-1^ will not be detected. This is an important result, because such enhanced transmission in tensed gels can significantly increase the effective range of communication between cells, pushing forward our understanding of processes such as cancer metastasis, blood vessel growth and morphogenesis.

## Acknowledgements

We are grateful to Haim Diamant, Yael Roichman and Yair Shokef for illuminating discussions. RS acknowledges support by the ISRAEL SCIENCE FOUNDATION (grant no. 1289/20), AL acknowledges support by the ISRAEL SCIENCE FOUNDATION (grant no. 1474/16), and the Israeli Centers for Research Excellence (grant no. 1902/12).

