## Supplementary material for "Probing Local Force Propagation in Tensed Fibrous Gels": Supplamentary Information

### Supplementary Information
#### Probing Local Force Propagation in Tensed Fibruos Gels

April 25, 2022

#### 1 Suitable theoretical expressions for translating beads in an isotropic gels

The displacement of a bead of radius a subject to a force $F_i$ within an elastic isotropic medium is given by [1]:

$$u_i^{(1)} = \frac{1}{6\pi aG}\frac{5-6\nu}{4(1-\nu)}F_i \tag{S1}$$

Where $G$ is the shear modulus and $\nu$ is Poisson's ratio. The displacement of a bead in position $r >> a$ with respect to the driven bead is [1]:

$$u_i^{(2)}(\vec{r}) = \frac{3a}{2(5-6\nu)|r|}[(3-4\nu)\delta_{ij} + \hat{r}_i\hat{r}_j]u_j^{(1)} \equiv \tilde{A}_{ij}(\vec{r})u_j^{(1)} \tag{S2}$$

In this study, we focus on the dependence of $A_{ij}$ on the distance r from the driven bead, rather than on its azimuthal dependence. We thus average $A_{ij}$ over all angles $\phi = tan^{-1}(y/x)$:

$$A_{ij}(r) \equiv < \tilde{A}_{ij}(\vec{r} >_\phi = \frac{1}{2\pi}\frac{3a}{2(5-6\nu)}|r|\int_0^{2\pi}\begin{bmatrix} 3-4\nu+cos^2(\phi) & cos(\phi)sin(\phi) \\ cos(\phi)sin(\phi) & 3-4\nu+sin^2(\phi)\end{bmatrix} d\phi \tag{S3}$$

$$A_{ij}(r) = \frac{3(7-8\nu)a}{4(5-6\nu)r}\delta_{ij} \tag{S4}$$

We note that $\nu$ in these expression denotes Poisson's ratio for the "bare" network, in the absence of solvent, and it may deviate from the incompressible value of 1/2 (For example, it can be calculated as 1/4 assuming network deformations are affine [2]). However, the deviation from Stokes' law for reasonable values of $\nu$. is around 10% [3], so it is not within our experimental accuracy to determine $\nu$. For convenience, we will assume $\nu$.=1/2, and it is to be noted that this assumption may introduce an error of up to 10% in our calculation of the shear modulus. Under this assumption, equation 1 yields the renowned generalized Stokes' law:

$$u_i^{(1)} = \frac{1}{6\pi aG}F_i \tag{S5}$$

And equation S4 yields:

$$A_{ij}(r) = \frac{9a}{8r}\delta_{ij} \tag{S6}$$

#### 2 Tensile Device

Figure S1 shows our simple tensile device with an attached silicone strip holding a fibrin gel, before and after applying tension. Ink lines on silicon strip were marked help determine the degree of stretch. The gels where tensed so that the diameter of the holes increased by 50%.

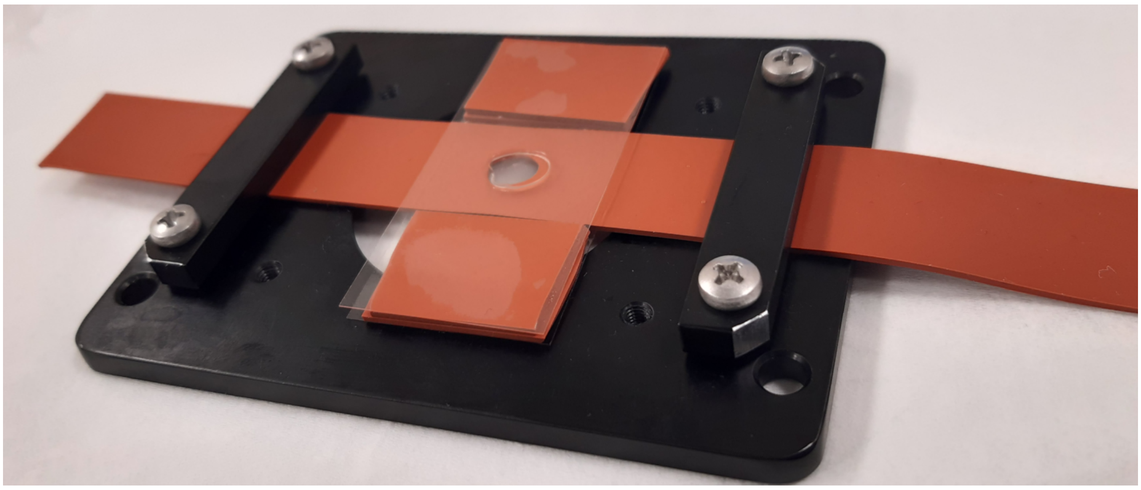

Figure S1: Tensile device used in the study. The long silicone strip, with a hole containing the fibrin gel, is gently placed on top of a glass slide on the device platform. The strip is clamped on one side, manually tensed to a pre-inscribed mark and clamped on the other side. A second glass slide is then placed above the gel, separated from it using spacers. The chamber is then closed from the sides with wet kim-wipes to maintain moisture.

### 3 Accuracy of bead amplitude measurement

To estimate the accuracy of our measurement of bead amplitudes, we perform experiments where we trap a 1.5 diameter bead in water. We oscillate the trap at varying trap amplitudes. We briefly demonstrate that under our experimental conditions, beads are expected to follow the trap with negligible damping due to water drag. The Fourier-space ensemble-averaged trajectory of a bead dispersed in water and driven by an oscillation optical trap is [4]:

$$< \tilde{x}_{bead}(\omega) >= \frac{\kappa \tilde{x}_{trap}(\omega)}{\kappa - m\omega^2 + i\omega\gamma} \tag{S8}$$

Where $\kappa$ is the trap stiffness, $m$ is the bead mass and $\gamma = 6\pi a\eta$ the drag coefficient. For our system,$\omega$ $1Hz$, $\kappa$ $10^{-4}N/m$, $m\omega^2$ $10^{-15}$ N/m and $\omega\gamma = 6\pi\omega a\eta$ $10^{-8}N/m$. It is therefore clear that the trap stiffness $\kappa$ strongly dominates the denominator and we can assume $< x_{bead} >= x_{trap}$ with negligible corrections. The contribution of thermal fluctuations can also be neglected on similar basis.

Figure S2 plots the bead amplitude measured from video tracking algorithm, vs the trap amplitude. It can be seen that the tracking algorithm correctly reproduces the trap amplitude even for amplitudes as small as 1 nm. These results demonstrate that the implemented tracking algorithm combined with careful Fourier analysis has a resolution of about 1/80 of the pixel size.

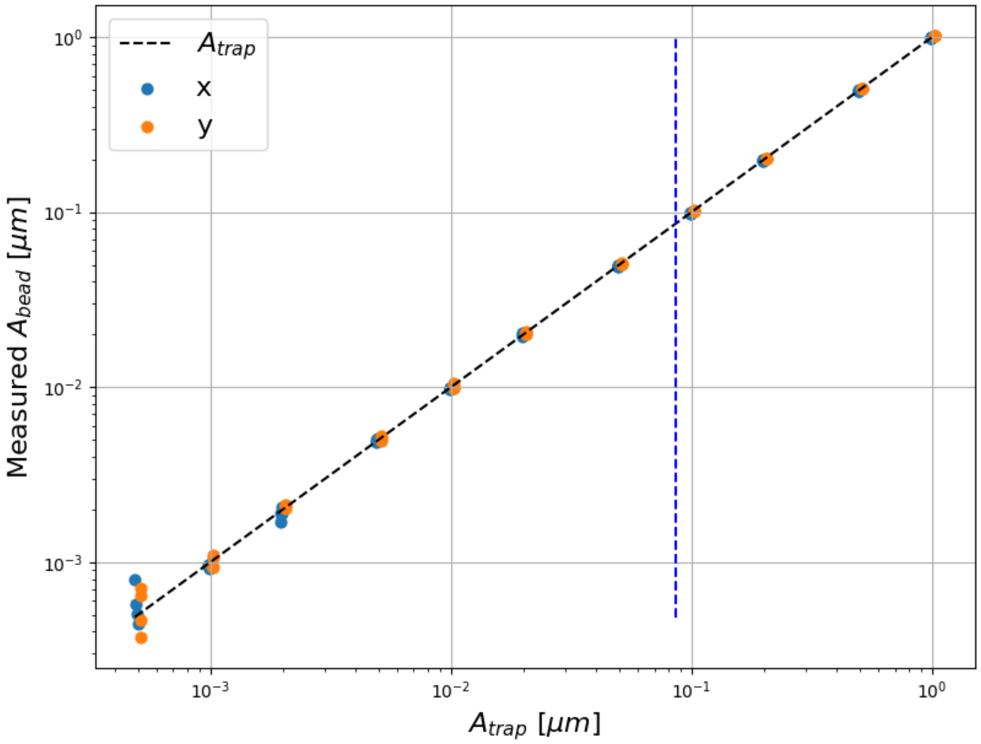


Figure S2: Estimating the accuracy of amplitude measurements. Obtained bead amplitudes match the programmed trap amplitudes down to 1nm oscillations. Blue dashed line indicates the pixel size.

### 4 Quantifying alignment in confocal images

Local image gradients are computed by a convolution of the skeletonized images with 21 by 21 Sobel operators [5], defined as:

$$h_x(i,j) = \frac{j}{i^2 + j^2}; h_y(i,j) = \frac{i}{i^2 + j^2} \tag{S7}$$

Where i and j are the row and columns and vary between -10 and 10. The x and y gradients $g_x$ and $g_y$ are computed by convolution of the images with $h_x$ and $h_y$ respectively. Local directionality $\theta = tan^{-1}(g_x/g_y)$ is computed and the nematic order paramete is computed as $NOP =< cos(2\theta) >$.

Most confocal scans were taken with x as the fast scanning axis. To remove directionality artifacts resulting from the fast axis choice in confocal microscopy, we recorded a subset of confocal scans with y as the fast scanning axis. Figure S3 shows the differences in the distribution of NOP values for scans with different fast axes. For this plot, only isotropic gel scans were included. It can be seen that scans in the x direction had an average NOP of 0.03, while scans in the y direction had the opposite value -0.03. We therefore concluded that for our quantification algorithm, the scan axis introduces a systematic error of 0.03 in the computation of the NOP, and we therefore subtract or add this value depending on the scan axis.

### 5 Verifying that material response is linear and frequency independent

We verified that the material response is linear by quantifying the response tensor $A_{ij}(r)$ for 3 different oscillation amplitudes. This was done for 2 untensed and 2 tensed gels. Since the quantification of $A_{ij}(r)$ includes normalizing by the driven bead amplitude, it's value should be

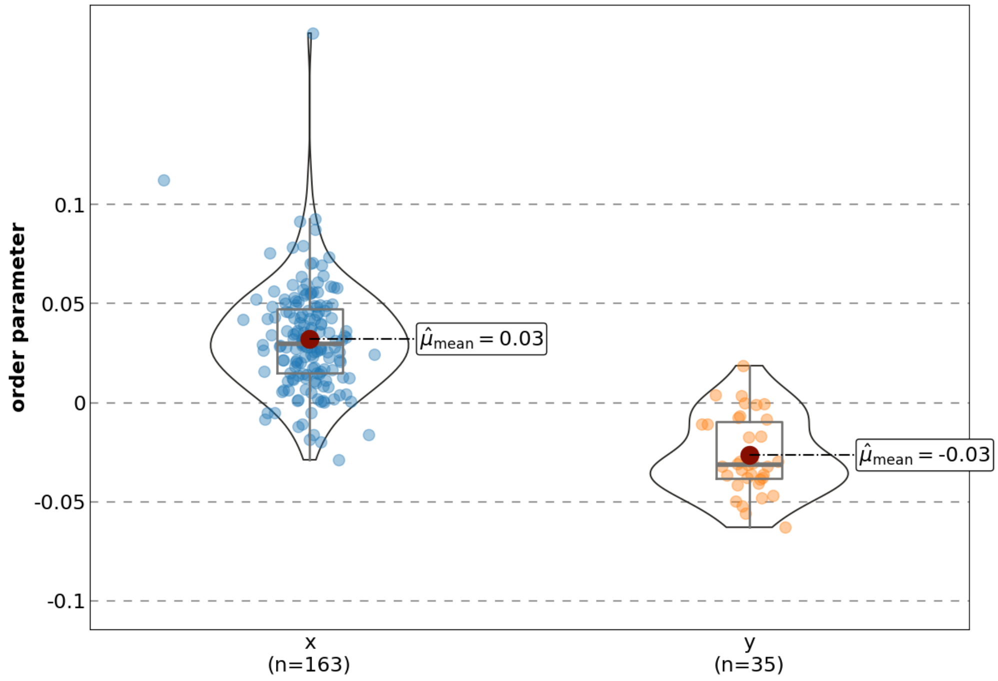


Figure S3: Compensating for alignment artifacts introduced by confocal scanning. When comparing the distribution of order parameter values for images scanned with x and y as the fast axis, we obtain a bias of 0.03 in the calculated order parameter, that is positive when scanning with x as the fast axis and negative when y is the fast axis. This bias is therefore subtracted from the computed NOP values in the main text.

identical for different oscillation amplitudes in the case of a linear response. In figure S4, the entries $A_{xx}$ and $A_{yy}$ of the response tensor are plotted as a function of normalized distance from the driven bead and for 3 different amplitude, in each of the gels. It can be seen that in all cases, the results are insensitive to the driven bead amplitude, as expected for a linear response.

In figure S5, we measure the 2-particle microrheology of an untensed gel at different frequencies. It can be seen that the response is not sensitive to frequency changes. Although we do expect to find a different response at frequencies higher than 10 Hz, we focus on the low frequency response, which is relevant for most biological processes. We show here that our chosen frequency of 0.5 Hz is well within the low-frequency plateau, as is also seen in bulk rheology and passive microrheology.

#### 6 Comparing individual gels

In Figures S6 and S7 we present the elements of the response tensor for 3 untensed and 3 tensed gels. We observe that the main features distinguishing untensed and tensed gels are robust: Isotropy of untensed gels, manifested by equal $A_x x$ and $A_y y$ curves, compared to unequal $A_x x$ and $A_y y$ for tensed gels. We also observe irregular power-law behavior in tensed gels, where the response decays slower than predicted by linear theory, as discussed in the main text.

#### 7 supplementary videos

Video 1 displays an oscillation of a bead in a labeled fibrin gel. The video was recorded by sequentially displacing the bead using an optical trap and taking a confocal scan of the surrounding gel. The fibers are shown in green and tracer beads are shown in orange.

Videos 2 and 3 display brightfield videos of a bead driven in the x and y directions, respectively. brightfield images were used in order to quantify the material response in practice.

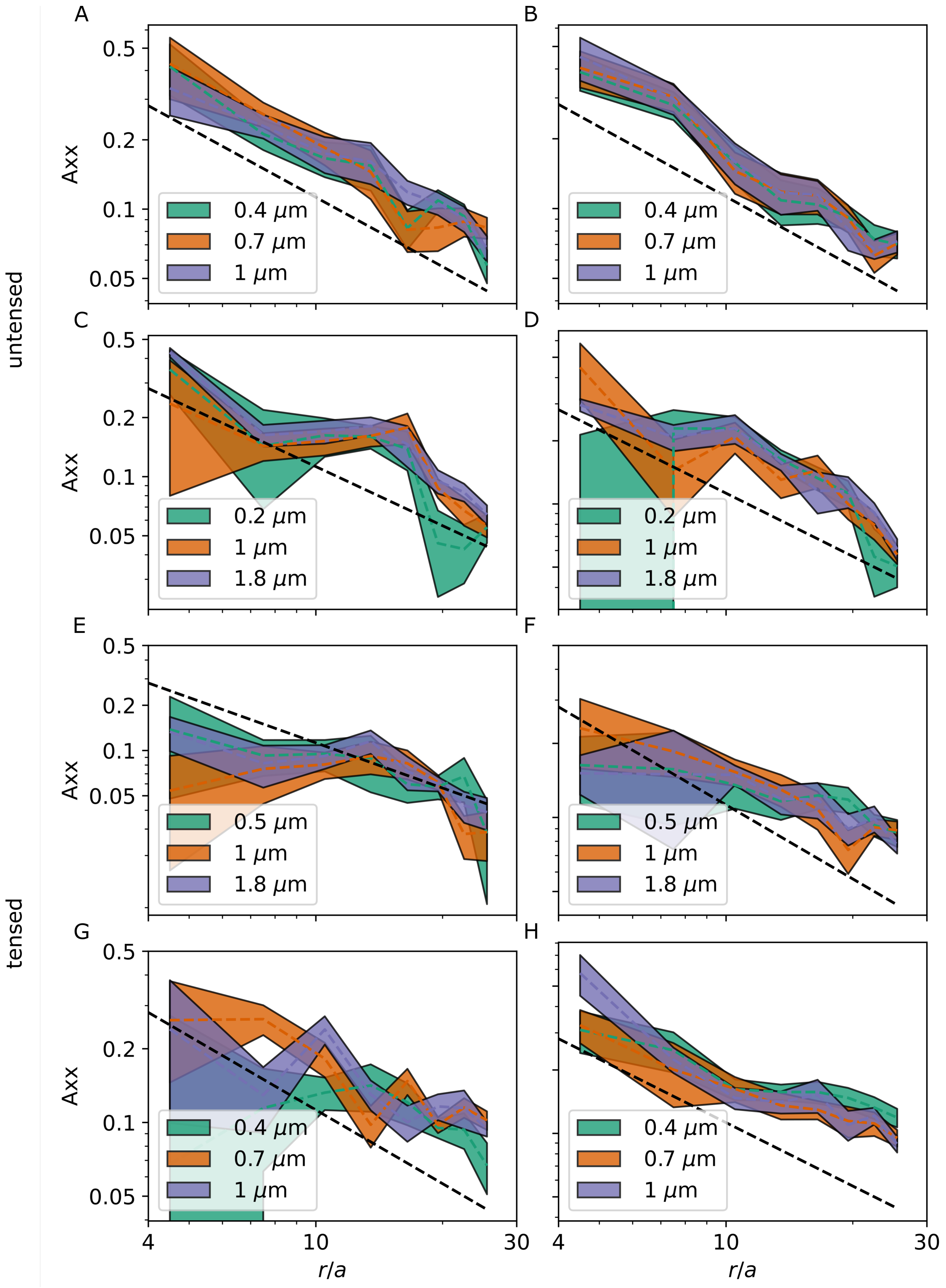


Figure S4: The response tensor elements at different amplitudes. (A),(B) and (C),(D) display the xx, yy elements for 2 different untensed gels, and (E),(F) and (G),(H) are for 2 different tensed gels. In all cases, it can be seen that the values measured are roughly the same for different oscillation amplitudes, indicating a linear response.

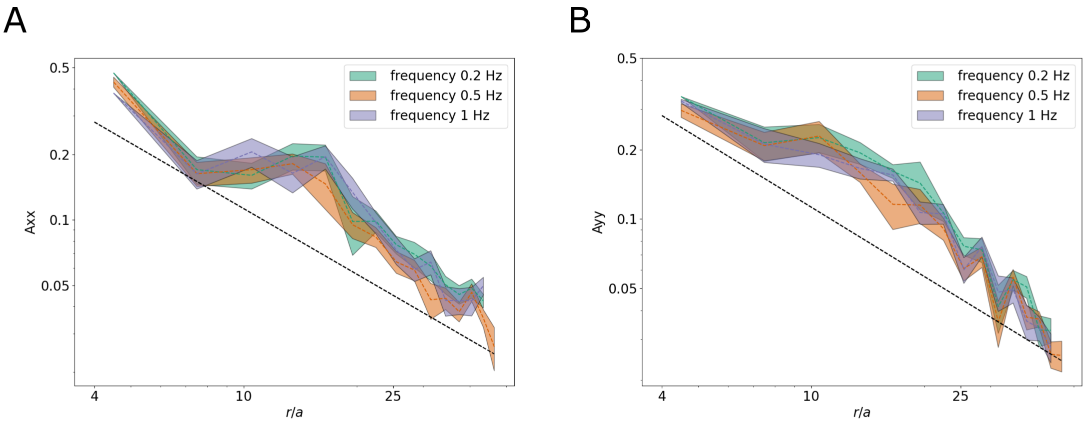


Figure S5: Frequency independence of the response tensor. The response tensor in an untensed gel was characterized, showing no significant dependence of frequency around the chosen frequency of 0.5Hz.

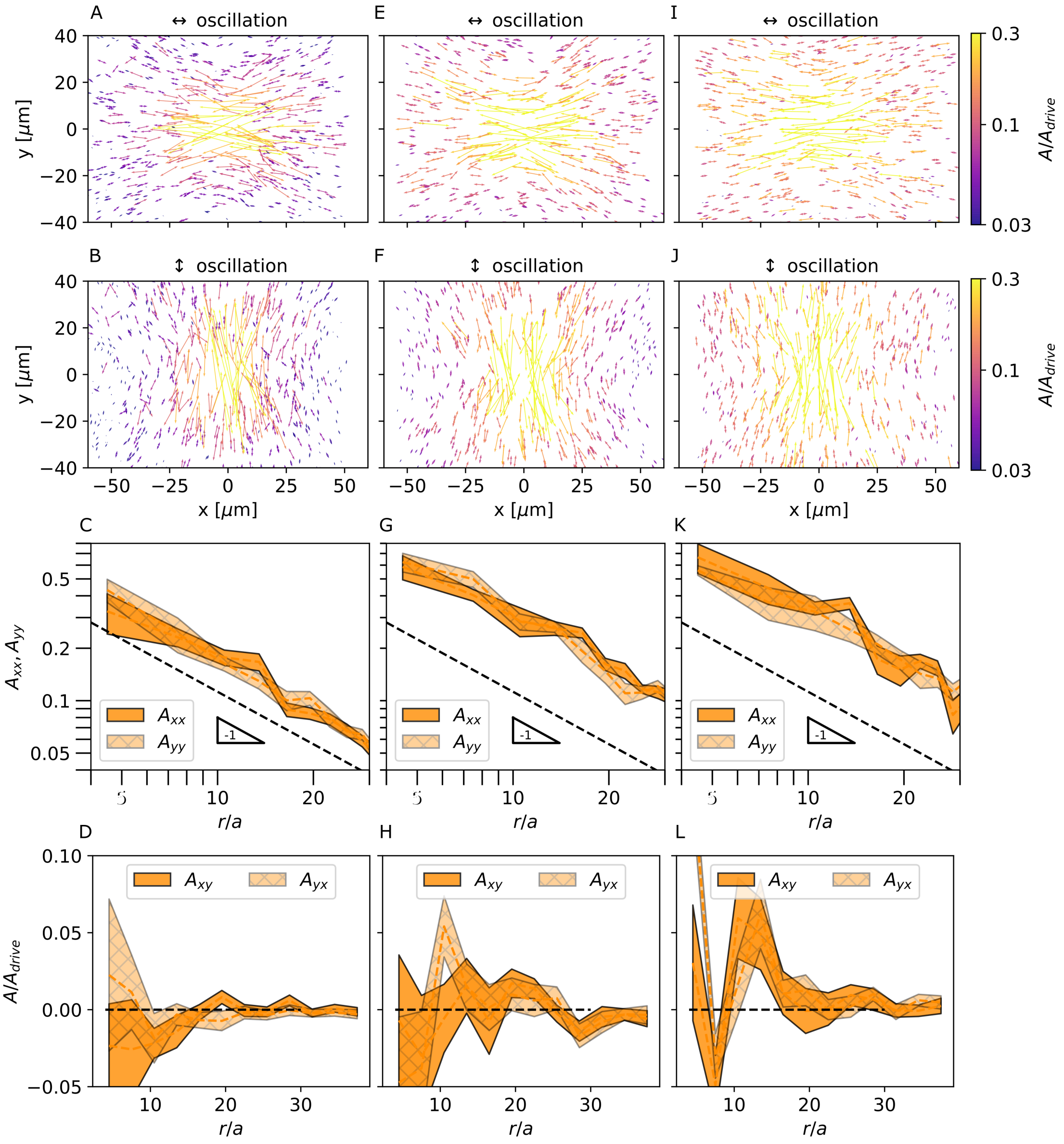


Figure S6: Comparing the measured response tensor for 3 individual untensed gels. Plots (A)-(D), (E)-(H) and (I)-(L) correspond to the 3 different gels, showing the robustness of the main properties of the response tensor discussed in the main text.

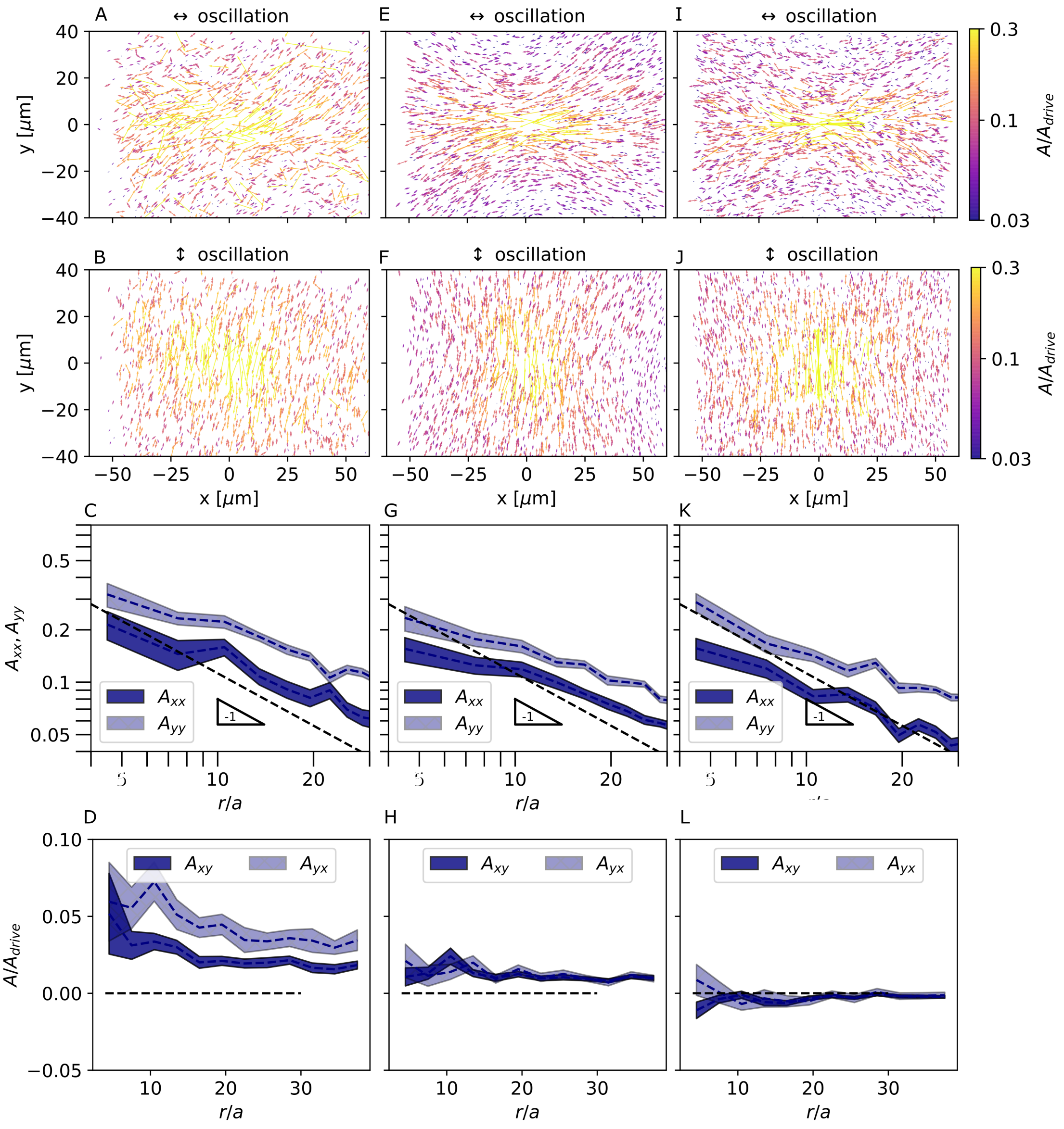


Figure S7: Comparing the measured response tensor for 3 individual tensed gels. Plots (A)-(D), (E)-(H) and (I)-(L) correspond to the 3 different gels, showing the robustness of the main properties of the response tensor discussed in the main text.
